# pcaExplorer: an R/Bioconductor package for interacting with RNA-seq principal components

**DOI:** 10.1101/493551

**Authors:** Federico Marini, Harald Binder

## Abstract

**Background:** Principal component analysis (PCA) is frequently useentirely written ind in genomics applications for quality assessment and exploratory analysis in high-dimensional data, such as RNA sequencing (RNA-seq) gene expression assays. Despite the availability of many software packages developed for this purpose, an interactive and comprehensive interface for performing these operations is lacking.

**Results:** We developed the `pcaExplorer` software package to enhance commonly performed analysis steps with an interactive and user-friendly application, which provides state saving as well as the automated creation of reproducible reports. `pcaExplorer` is implemented in R using the Shiny framework and exploits data structures from the open-source Bioconductor project. Users can easily generate a wide variety of publication-ready graphs, while assessing the expression data in the different modules available, including a general overview, dimension reduction on samples and genes, as well as functional interpretation of the principal components.

**Conclusion:** `pcaExplorer` is distributed as an R package in the Bioconductor project (http://bioconductor.org/packages/pcaExplorer/), and is designed to assist a broad range of researchers in the critical step of interactive data exploration.

## Background

Transcriptomic data via RNA sequencing (RNA-seq) aim to measure gene/transcript expression levels, summarized from the tens of millions of reads generated by next generation sequencing technologies (Mortazavi *et al.*, 2008). Besides standardized workflows and approaches for statistical testing, tools for exploratory analysis of such large data volumes are needed. In particular, after counting the number of reads that overlap annotated genes, using tools such as featureCounts (Liao *et al.*, 2014) or HTSeq (Anders *et al.*, 2015), the result still is a high-dimensional matrix of the transcriptome profiles, with rows representing features (e.g., genes) and columns representing samples (i.e. the experimental units). This matrix constitutes an essential intermediate result in the whole process of analysis (Anders *et al.*, 2013; Conesa *et al.*, 2016), irrespective of the specific aim of the project.

A wide number and variety of software packages have been developed for accommodating the needs of the researcher, mostly in the R/Bioconductor framework (Gentleman *et al.*, 2004; Huber *et al.*, 2015). Many of them focus on the identification of differentially expressed genes (Love *et al.*, 2014; McCarthy *et al.*, 2012) for discovering quantitative changes between experimental groups, while others address alternative splicing, discovery of novel transcripts or RNA editing.

Exploratory data analysis is a common step to all these workflows (Conesa *et al.*, 2016), and constitutes a key aspect for the understanding of complex biological systems, by indicating potential problems with the data and sometimes also for generating new hypotheses. Despite its importance for generating reliable results, e.g. by helping the researchers uncovering outlying samples, or diagnosing batch effects, this analysis workflow component is often neglected, as many of the steps involved might require a considerable proficiency of the user in the programming languages.

Among the many techniques adopted for exploring multivariate data like transcriptomes, principal component analysis (PCA, (Jolliffe, 2002)) is often used to obtain an overview of the data in a low-dimensional subspace (Yeung and Ruzzo, 2001; Ma and Dai, 2011). Implementations where PCA results can be explored are available, mostly focused on small sample datasets, such as Fisher’s iris (Fisher, 1984) (https://gist.github.com/dgrapov/5846650 or https://github.com/dgrapov/DeviumWeb, https://github.com/benmarwick/Interactive_PCA_Explorer) and have been developed rather for generic data, without considering the aspects typical of transcriptomic data (http://langtest.jp/shiny/pca/, (Vaissie *et al.*, 2017)). In the field of genomics, some tools are already available for performing such operations (Sharov *et al.*, 2005; la Grange *et al.*, 2009; Metsalu and Vilo, 2015; Khomtchouk *et al.*, 2016; Manimaran *et al.*, 2016; Nelson *et al.*, 2016; Schultheis *et al.*, 2018), yet none of them feature an interactive analysis, fully integrated in Bioconductor, while also providing the basis for generating a reproducible analysis (Peng, 2011; McNutt, 2014). Alternatively, more general software suites are also available (e.g. Orange, https://orange.biolab.si), designed as user interfaces offering a range of data visualization, exploration, and modeling techniques.

Our solution, `pcaExplorer`, is a web application developed in the Shiny framework (Chang *et al.*, 2018), which allows the user to efficiently explore and visualize the wealth of information contained in RNA-seq datasets with PCA, performed for visualizing relationships either among samples or genes. `pcaExplorer` additionally provides other tools typically needed during exploratory data analysis, including normalization, heatmaps, boxplots of shortlisted genes and functional interpretation of the principal components. We included a number of coloring and customization options to generate and export publication-ready vector graphics.

To support the reproducible research paradigm, we provide state saving and a text editor in the app that fetches the live state of data and input parameters, and automatically generates a complete HTML report, using the `rmarkdown` and `knitr` packages (Allaire *et al.*, 2018; Xie, 2015), which can e.g. be readily shared with collaborators.

## Implementation

### General design of pcaExplorer

`pcaExplorer` is entirely written in the R programming language and relies on several other widely used R packages available from Bioconductor. The main functionality can be accessed by a single call to the `pcaExplorer`() function, which starts the web application.

The interface layout is built using the `shinydashboard` package (Chang and Borges Ribeiro, 2018), with the main panel structured in different tabs, corresponding to the dedicated functionality. The sidebar of the dashboard contains a number of widgets which control the app behavior, shared among the tabs, regarding how the results of PCA can be displayed and exported. A task menu, located in the dashboard header, contains buttons for state saving, either as binary RData objects, or as environments accessible once the application has been closed.

A set of tooltips, based on bootstrap components in the `shinyBS` package (Bailey, 2015), is provided throughout the app, guiding the user for choosing appropriate parameters, especially during the first runs to get familiar with the user interface components. Conditional panels are used to highlight which actions need to be undertaken to use the respective tabs (e.g., principal components are not computed if no normalization and data transformation have been applied).

Static visualizations are generated exploiting the base and `ggplot2` (Wickham, 2016) graphics systems in R, and the possibility to interact with them (zooming in and displaying additional annotation) is implemented with the rectangular brushing available in the Shiny framework. Moreover, fully interactive plots are based on the `d3heatmap` and the `threejs` packages (Cheng and Galili, 2018; Lewis, 2017). Tables are also displayed as interactive objects for easier navigation, thanks to the DT package (Xie, 2018).

The combination of `knitr` and R Markdown allows to generate interactive HTML reports, which can be browsed at runtime and subsequently exported, stored, or shared with collaborators. A template with a complete analysis, mirroring the content of the main tabs, is provided alongside the package, and users can customize it by adding or editing the content in the embedded editor based on the `shinyAce` package (Nijs *et al.*, 2018).

`pcaExplorer` has been tested on macOS, Linux, and Windows. It can be downloaded from the Bioconductor project page (http://bioconductor.org/packages/pcaExplorer/), and its development version can be found at https://github.com/federicomarini/pcaExplorer/. Moreover, `pcaExplorer` is also available as a Bioconda recipe (Grüning *et al.*, 2018), to make the installation procedure less complicated (binaries at https://anaconda.org/bioconda/bioconductor-pcaExplorer), as well to provide the package in isolated software environments, reducing the burden of software version management.

A typical modern laptop or workstation with at least 8 GB RAM is sufficient to run `pcaExplorer` on a variety of datasets. While the loading and preprocessing steps can vary according to the dataset size, the time required for completing a session with `pcaExplorer` mainly depends on the depth of the exploration. We anticipate a typical session could take approximately 15-30 minutes (including the report generation), once the user has familiarized with the package and its interface.

### Typical usage workflow

Figure 1 illustrates a typical workflow for the analysis with `pcaExplorer`. `pcaExplorer` requires as input two fundamental pieces of information, i.e. the raw count matrix, generated after assigning reads to features such as genes via tools such as HTSeq-count or featureCounts, and the experimental metadata table, which contains the essential variables for the samples of interest (e.g., condition, tissue, cell line, sequencing run, batch, library type, …). The information stored in the metadata table is commonly required when submitting the data to sequencing data repositories such as NCBI’s Gene Expression Omnibus (https://www.ncbi.nlm.nih.gov/geo/), and follows the standard proposed by the FAIR Guiding Principles (Wilkinson *et al.*, 2016).

**Figure 1:**
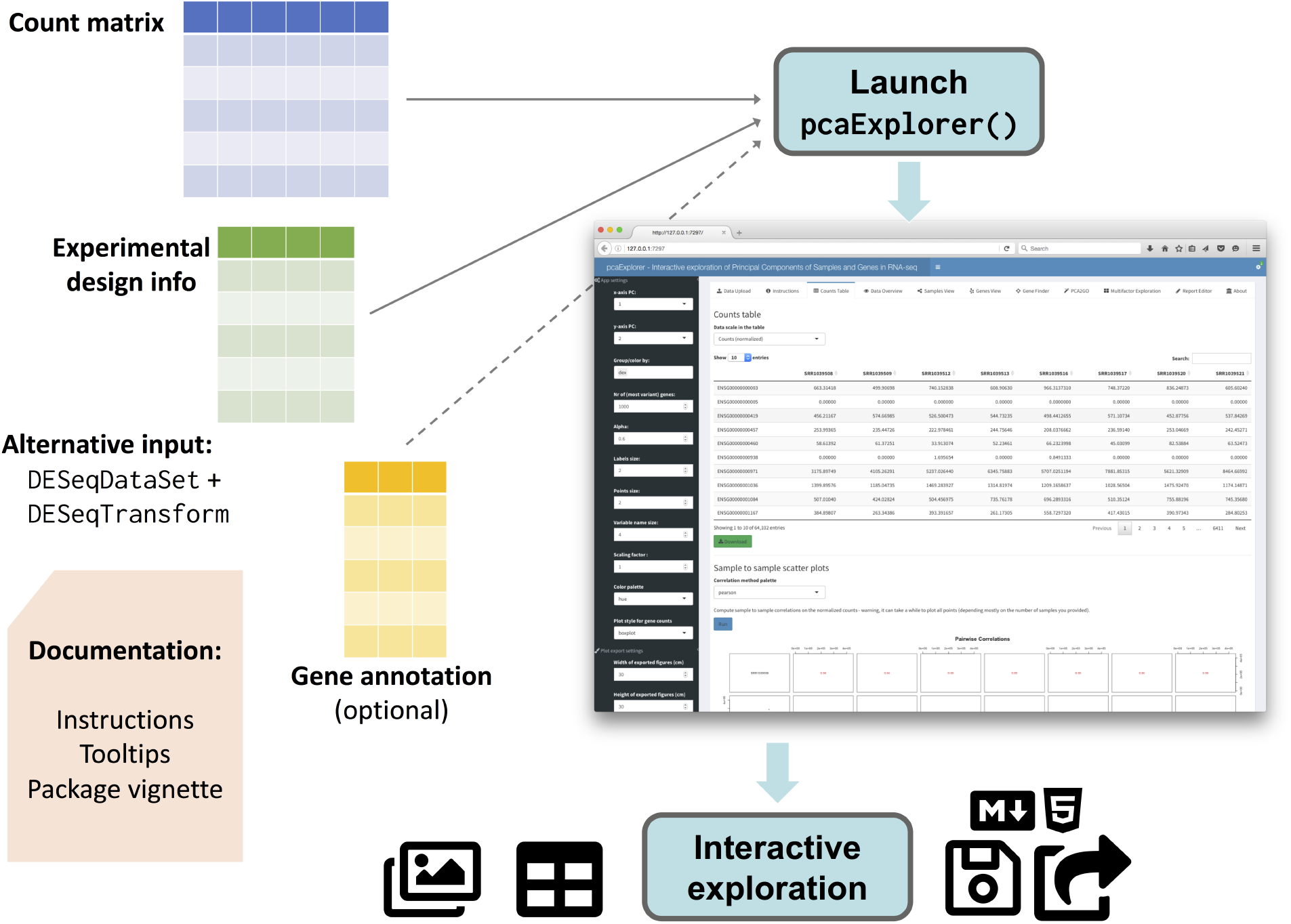
Overview of the pcaExplorer workflow. A typical analysis with `pcaExplorer` starts by providing the matrix of raw counts for the sequenced samples, together with the corresponding experimental design information. Alternatively, a combination of a `DESeqDataSet` and a `DESeqTransform` objects can be given as input. Specifying a gene annotation can allow displaying of alternative IDs, mapped to the row names of the main expression matrix. Documentation is provided at multiple levels (tooltips and instructions in the app, on top of the package vignette). After launching the app, the interactive session allows detailed exploration capability, and the output can be exported (images, tables) also in form of a R Markdown/HTML report, which can be stored or shared. (Icons contained in this figure are contained in the collections released by Font Awesome under the CC BY 4.0 license.)

The count matrix and the metadata table can be provided as parameters by reading in delimiter-separated (tab, comma, or semicolon) text files, with identifiers as row names and a header indicating the ID of the sample, or directly uploaded while running the app. A preview of the data is displayed below the widgets in the *Data Upload* tab, as an additional check for the input procedures. Alternatively, this information can be passed in a single object, namely a `DESeqDataSet` object, derived from the broadly used `SummarizedExperiment` class (Huber *et al.*, 2015). The required steps for normalization and transformation are taken care of during the preprocessing phase, or can be performed in advance. If not specified when launching the application, `pcaExplorer` automatically computes normalization factors using the `estimateSizeFactors`() function in the `DESeq2` package, which has been shown to perform robustly in many scenarios under the assumption that most of the genes are not differentially expressed (Dillies *et al.*, 2013).

Two additional objects can be provided to the `pcaExplorer`() function: the annotation object is a data frame containing matched identifiers for the features of interest, encoded with different key types (e.g., ENTREZ, ENSEMBL, HGNC-based gene symbols), and a pca2go object, structured as a list containing enriched GO terms (Ashburner *et al.*, 2000) for genes with high loadings, in each principal component and in each direction. These elements can also be conveniently uploaded or calculated on the fly, and make visualizations and insights easier to read and interpret.

Users can resort to different venues for accessing the package documentation, with the vignette also embedded in the web app, and the tooltips to guide the first steps through the different components and procedures.

Once the data exploration is complete, the user can store the content of the reactive values in binary RData objects, or as environments in the R session. Moreover, all available plots and tables can be manually exported with simple mouse clicks. The generation of an interactive HTML report can be meaningfully considered as the concluding step. Users can extend and edit the provided template, which seamlessly retrieves the values of the reactive objects, and inserts them in the context of a literate programming compendium (Knuth, 1984), where narrated text, code, and results are intermixed together, providing a solid means to warrant the technical reproducibility of the performed operations.

### Deploying pcaExplorer on a Shiny server

In addition to local installation, `pcaExplorer` can also be deployed as a web application on a Shiny server, such that users can explore their data without the need of any extra software installation. Typical cases for this include providing a running instance for serving members of the same research group, setup by a bioinformatician or a IT-system admin, or also allowing exploration and showcasing relevant features of a dataset of interest.

A publicly available instance is accessible at http://shiny.imbei.uni-mainz.de:3838/pcaExplorer, for demonstration purposes, featuring the primary human airway smooth muscle cell lines dataset (Himes *et al.*, 2014). To illustrate the full procedure to setup `pcaExplorer` on a server, we documented all the steps at the GitHub repository https://github.com/federicomarini/pcaExplorer_serveredition. Compared to web services, our Shiny app (and server) approach also allows for protected deployment inside institutional firewalls to control sensitive data access.

### Documentation

The functionality indicated above and additional functions, included in the package for enhancing the data exploration, are comprehensively described in the package vignettes, which are also embedded in the Instructions tab.

Extensive documentation for each function is provided, and this can also be browsed at https://federicomarini.github.io/pcaExplorer/, built with the `pkgdown` package (Wickham and Hesselberth, 2018). Notably, a dedicated vignette describes the complete use case on the `airway` dataset, and is designed to welcome new users in their first experiences with the `pcaExplorer` package (available at http://federicomarini.github.io/pcaExplorer/articles/upandrunning.html).

## Results

### Data input and overview

Irrespective of the input modality, two objects are used to store the essential data, namely a `DESeqDataSet` and a `DESeqTransform`, both used in the workflow based on the `DESeq2` package (Anders *et al.*, 2013). Different data transformations can be applied in `pcaExplorer`, intended to reduce the mean-variance dependency in the transcriptome dataset: in addition to the simple shifted log transformation (using small positive pseudocounts), it is possible to apply a variance stabilizing transformation or also a regularized-logarithm transformation. The latter two approaches help for reducing heteroscedasticity, to make the data more usable for computing relationships and distances between samples, as well as for visualization purposes (Love *et al.*, 2015).

The data tables for raw, normalized (using the median of ratios method in DESeq2), and transformed data can be accessed as interactive table in the *Counts Table* module. A scatter plot matrix for the normalized counts can be generated with the matrix of the correlation among samples.

Further general information on the dataset is provided in the *Data Overview* tab, with summaries over the design metadata, library sizes, and an overview on the number of robustly detected genes. Heatmaps display the distance relationships between samples, and can be decorated with annotations based on the experimental factors, selected from the sidebar menu. Fine-grained control on all the downstream operations is provided by the series of widgets located on the left side of the app. These include, for example, the number of most variant genes to include for the downstream steps, as well as graphical options for tailoring the plots to export them ready for publication.

### Exploring Principal Components

The *Samples View* tab (Figure 2A) provides a PCA-based visualization of the samples, which can be plotted in 2 and 3 dimensions on any combination of PCs, zoomed and inspected, e.g. for facilitating outlier identification. A scree plot, helpful for selecting the number of relevant principal components, and a plot of the genes with highest loadings are also given in this tab. The *Genes View* tab, displayed in Figure 2B, is based on a PCA for visualizing a user-defined subset of most variant genes, e.g. to assist in the exploration of potentially interesting clusters. The samples information is combined in a biplot for better identification of PC subspaces. When selecting a region of the plot and zooming in, heatmaps (both static and interactive) and a profile plot of the corresponding gene subset are generated. Single genes can also be inspected by interacting with their names in the plot. The underlying data, displayed in collapsible elements to avoid cluttering the user interface, can also be exported in tabular text format.

**Figure 2:**
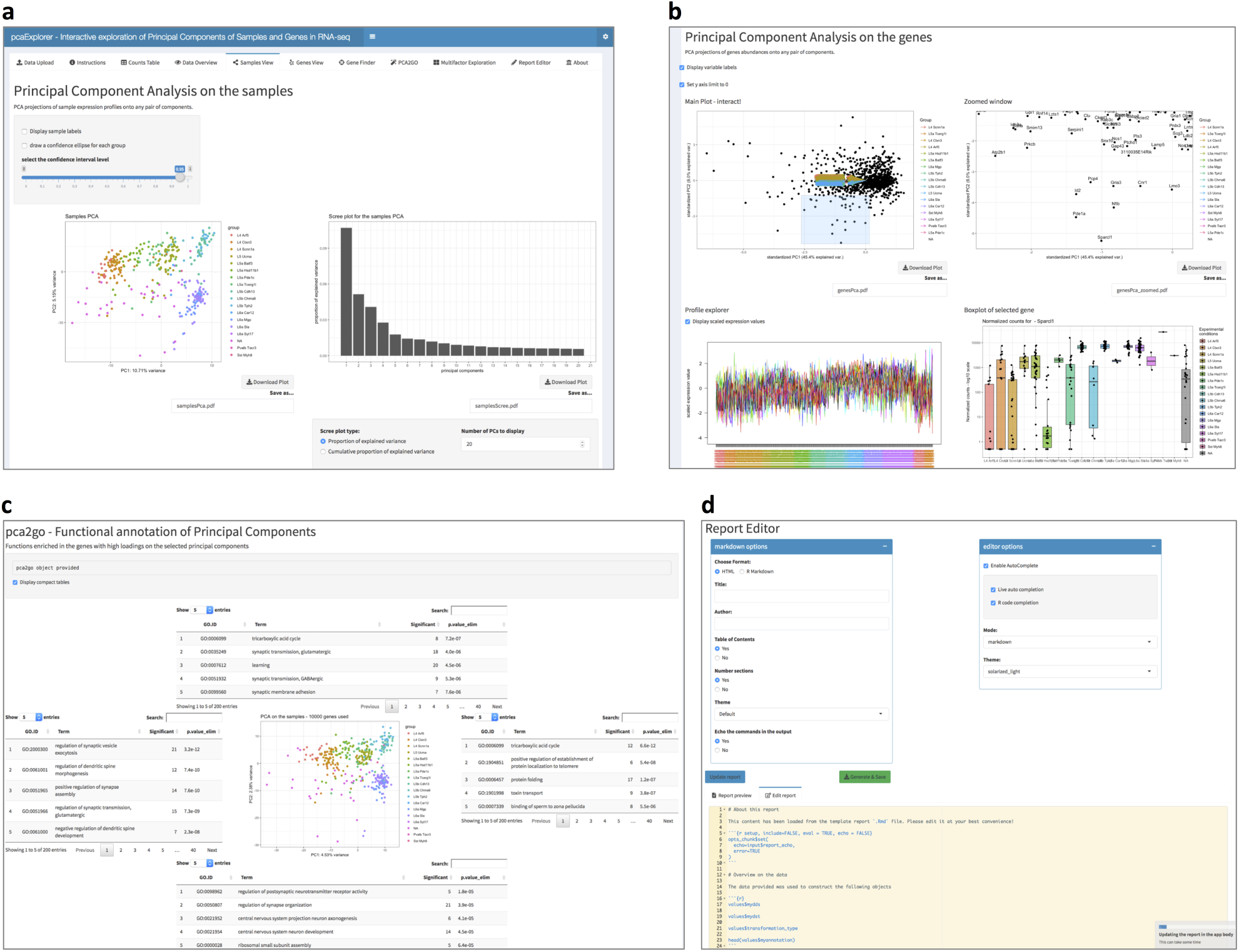
Selected screenshots of the pcaExplorer application. **a** Principal components from the point of view of the samples, with a zoomable 2D PCA plot (3D now shown due to space) and a scree plot. Additional boxes show loadings plots for the PCs under inspection, and let users explore the effect of the removal of outlier samples. **b** Principal components, focused on the gene level. Genes are shown in the PCA plot, with sample labels displayed as in a biplot. A profile explorer and heatmaps (not shown due to space) can be plotted for the subset selected after user interaction. Single genes can also be inspected with boxplots. **c** Functional annotation of principal components, with an overview of the GO-based functions enriched in the loadings in each direction for the selected PCs. The pca2go object can be provided at launch, or also computed during the exploration. **d** Report Editor panel, with markdown-related and general options shown. Below, the text editor displays the content of the analysis for building the report, defaulting to a comprehensive template provided with the package.

### Functional annotation of Principal Components

Users might be interested in enriching PCA plots with functional interpretation of the PC axes and directions. The *PCA2GO* tab provides such a functionality, based on the Gene Ontology database. It does so by considering subsets of genes with high loadings, for each PC and in each direction, in an approach similar to `pcaGoPromoter` (Hansen *et al.*, 2012). The functional categories can be extracted with the functions in `pcaExplorer` (`pca2go`() and `limmaquickpca2go()`), which conveniently wrap the implementation of the methods in (Alexa *et al.*, 2006; Ritchie *et al.*, 2015). This annotation is displayed in interactive tables which decorate a PCA plot, positioned in the center of the tab.

An example of this is shown in Figure 2C, where we illustrate the functionality of `pcaExplorer` on a single-cell RNA-seq dataset. This dataset contains 379 cells from the mouse visual cortex, and is a subset of the data presented in (Tasic *et al.*, 2016), included in the scRNAseq package (http://bioconductor.org/packages/scRNAseq/).

### Further data exploration

Further investigation will typically require a more detailed look at single genes. This is provided by the *Gene Finder* tab, which provides boxplots (or violin plots) for their distribution, superimposed by jittered individual data points. The data can be grouped by any combination of experimental factors, which also automatically drive the color scheme in each of the visualizations. The plots can be downloaded during the live session, and this functionality extends to the other tabs.

In the *Multifactor Exploration* tab, two experimental factors can be incorporated at the same time into a PCA visualization. As in the other PCA-based plots, the user can zoom into the plot and retrieve the underlying genes to further inspect PC subspaces and the identified gene clusters of interest.

### Generating reproducible results

The *Report Editor* tab (Figure 2D) provides tools for enabling reproducible research in the exploratory analysis described above. Specifically, this tab captures the current state of the ongoing analysis session, and combines it with the content of a pre-defined analysis template. The output is an interactive HTML report, which can be previewed in the app, and subsequently exported.

Experienced users can add code for additional analyses using the text editor, which supports R code completion, delivering an experience similar to development environments such as RStudio. Source code and output can be retrieved, combined with the state saving functionality (accessible from the app task menu), either as binary data or as object in the global R environment, thus guaranteeing fully reproducible exploratory data analyses.

## Discussion

The application and approach proposed by our package `pcaExplorer` aims to provide a combination of usability and reproducibility for interpreting results of principal component analysis and beyond.

Compared to the other existing software packages for genomics applications, `pcaExplorer` is released as a standalone package in the Bioconductor project, thus guaranteeing the integration in a system with daily builds which continuously check the interoperability with the other dependencies. Moreover, `pcaExplorer` fully leverages existing efficient data structures for storing genomic datasets (`SummarizedExperiment` and its derivatives), represented as annotated data matrices. Some applications (clustVis, START App, Wilson) are also available as R packages (either on CRAN or on GitHub), while others are only released as open-source repositories to be cloned (MicroScope).

Additionally, `pcaExplorer` can be installed both on a local computer, and on a Shiny server. This is particularly convenient when the application is to be accessed as a local instance by multiple users, as it can be the case in many research laboratories, working with unpublished or sensitive patient-related data. We provide extensive documentation for all the use cases mentioned above.

The functionality of `pcaExplorer` to deliver a template report, automatically compiled upon the operations and edits during the live session, provides the basis for guaranteeing the technical reproducibility of the results, together with the exporting of workspaces as binary objects. This aspect has been somewhat neglected by many of the available software packages; out of the ones mentioned here, BatchQC supports the batch compilation of a report based on the functions inside the package itself. Orange (https://orange.biolab.si) also allows the creation of a report with the visualizations and output generated at runtime, but this cannot be extended with custom operations defined by the user, likely due to the general scope of the toolbox.

Future work will include the exploration of other dimension reduction techniques (e.g. sparse PCA (Witten *et al.*, 2009) and t-SNE (van der Maaten and Hinton, 2008) to name a few), which are also commonly used in genomics applications, especially for single-cell RNA-seq data. The former method enforces the sparsity constraint on the input variables, thus making their linear combination easier to interpret, while t-SNE is a non-linear kernel-based approach, which better preserves the local structure of the input data, yet with higher computational cost and a non-deterministic output, which might be not convenient to calculate at runtime on larger datasets. For the analysis of single-cell datasets, additional preprocessing steps need to be taken before they can be further investigated with `pcaExplorer`. The results of these and other algorithms can be accommodated in Bioconductor containers, as proposed by the `SingleCellExperiment` class (as annotated `colData` and `rowData` objects, or storing low-dimensional spaces as slots of the original object), allowing for efficient and robust interactions and visualizations, e.g. side-by-side comparisons of different reduced dimension views.

## Conclusion

Here we presented `pcaExplorer`, an R/Bioconductor package which provides a Shiny web based interface for the interactive and reproducible exploration of RNA-seq data, with a focus on principal component analysis. It allows to perform the essential steps in the exploratory data analysis workflow in a user-friendly manner, displaying a variety of graphs and tables, which can be readily exported. By accessing the reactive values in the latest state of the application, it can additionally generate a report, which can be edited, reproduced, and shared among researchers.

As exploratory analyses can play an important role in many stages of RNA-seq workflows, we anticipate that `pcaExplorer` will be very generally useful, making exploration and other stages of genomics data analysis transparent and accessible to a broader range of scientists.

In summary, our package `pcaExplorer` aims to become a companion tool for many RNA-seq analyses, assists the user in performing a fully interactive yet reproducible exploratory data analysis, and is seamlessly integrated into the ecosystem provided by the Bioconductor project.

## Additional information

### Availability and requirements

**Project name:** pcaExplorer

**Project home page:** http://bioconductor.org/packages/pcaExplorer/ (release), https://github.com/federicomarini/pcaExplorer/ (development version)

**Archived version:** https://doi.org/10.5281/zenodo.2633159, package source as gzipped tar archive of the version reported in this article

**Project documentation:** rendered at https://federicomarini.github.io/pcaExplorer/

**Operating systems:** Linux, Mac OS, Windows

**Programming language:** R

**Other requirements:** R 3.3 or higher, Bioconductor 3.3 or higher

**License:** MIT

**Any restrictions to use by non-academics:** none.

## Abbreviations

CRAN: Comprehensive R Archive Network
GO: Gene Ontology
PC: Principal Component
PCA: Principal Component Analysis
RNA-seq: RNA sequencing
t-SNE: t-distributed Stochastic Neighbor Embedding

## Acknowledgements

We thank Sebastian Schubert and Carina Santos of the Ruf lab (CTH Mainz) for fruitful discussions and their feedback as early adopters of the `pcaExplorer` package, as well as the users’ community for their helpful suggestions. We also thank Miguel Andrade, Wolfram Ruf, Franziska Härtner, and Gerrit Toenges for their helpful comments on the manuscript.

## Availability of data and materials

Data used in the described use cases is available from the following articles:

- The airway smooth muscle cell RNA-seq is included in PubMed ID: 24926665. GEO entry: GSE52778, accessed from the Bioconductor experiment package `airway` (http://bioconductor.org/packages/airway/, version 0.114.0).
- The allen data set on single cell from from the mouse visual cortex is included in PubMed ID: 26727548. Accessed from the Bioconductor experiment package `scRNAseq` package(http://bioconductor.org/packages/scRNAseq/, version 1.6.0)

The `pcaExplorer` package can be downloaded from its Bioconductor page (http://bioconductor.org/packages/pcaExplorer/) or the GitHub development page (https://github.com/federicomarini/pcaExplorer/). `pcaExplorer` is also provided as a recipe in Bioconda (https://anaconda.org/bioconda/bioconductor-pcaExplorer).

## Authors’ contributions

FM conceived and implemented the `pcaExplorer` package, and wrote the manuscript. HB supervised the implementation and edited the manuscript. Both authors read and approved the final version of the manuscript.

## Ethics approval and consent to participate

Not applicable.

## Consent for publication

Not applicable.

